# L-ascorbic acid (vitamin C) fermentation by the human pathogen *Vibrio cholerae*

**DOI:** 10.1101/2020.09.08.288738

**Authors:** J.R. Rosenberger, N.D. McDonald, E.F. Boyd

**Affiliations:** Department of Biological Sciences, University of Delaware, Newark, DE, 19716

## Abstract

L-ascorbic acid, commonly known as vitamin C, is a ubiquitous 6-carbon carbohydrate characterized by its ability to scavenge free radicals. In enteric bacteria, L-ascorbate can be utilized as a nutrient using the UlaABCDEF and UlaG-UlaRpathway under anaerobic conditions. In this study, we identified homologs of the Ula system within *Vibrio cholerae* and showed that *V. cholerae* is able to utilize L-ascorbate as an energy source. Growth pattern assays of a *ulaG* in-frame deletion mutant demonstrated that *ulaG* is essential for L-ascorbate fermentation. Expression analysis showed that *ula* catabolism and transport genes were significantly induced in cells grown in the presence of L-ascorbate compared to glucose and these genes were also highly induced during growth on intestinal mucus. In *in vitro* growth competition assays, the *ulaG* mutant was outcompeted by wild type when grown in intestinal mucus suggesting the Ula system could be important for fitness. Within the *ula* operon in *V. cholerae* and all *Vibrio* species a homology of ORF VCA0243 is present that encodes a pyridoxal phosphate (PLP) phosphatase. This enzyme in *E. coli*, converts the active form of vitamin B_6_ PLP to its inactive form pyridoxal (PL). In *V. splendidus* and related species, the aerobic and anaerobic L-ascorbate pathway genes cluster together and both systems contain a PLP phosphatase. An in-frame deletion mutant of *vca0243* resulted in a growth defect in L-ascorbate fermentation as well as additional carbon and amino acid sources indicating a role in cellular metabolism. Phylogenetic analysis of UlaG and UlaD suggested the region was acquired by horizontal gene transfer.

**Importance:** L-ascorbate is a carbohydrate present in the human intestine, available for microbial consumption and several enteric species have been shown to utilize this compound as an energy source. We demonstrated that L-ascorbate fermentation genes are also present among marine bacteria from the family *Vibrionaceae* and that the human pathogen *V. cholerae* can ferment L-ascorbate as an energy source. Within the Ula operon in all *Vibrionaceae*, a putative pyridoxal phosphate phosphatase was present that was required for L-ascorbate fermentation and cellular metabolism in general. The Ula system was present among a limited number of genera within *Vibrionaceae; Vibrio, Aliivibrio* and *Photobacterium* and showed an evolutionary history consistent with horizontal transfer between genera and species.

## Introduction

L-Ascorbic acid is a ubiquitous six-carbon carbohydrate known for its role as an antioxidant. It is also an essential nutrient, found within GI tissues, gastric juices as well as the mucosa, required for enzyme function and tissue repair in humans and other animals (1-3). Some higher order eukaryotes, such as humans, have lost the ability to biosynthesize ascorbate (4-6). In these species, the requirement for ascorbate is met through dietary intake(4, 5).

It is known that enteric bacteria are capable of utilizing L-ascorbate as a carbon and energy source (7-12). In *E. coli*, two systems exist for the catabolism of L-ascorbate, the Ula system and the YiaK-YiaS system. The Ula system is required for the fermentation of L-ascorbate under anaerobic conditions, whereas under aerobic conditions, both the Ula system and the YiaK-YiaS system are required. Studies in *E. coli* and *Klebsiella pneumoniae* show neither can aerobically catabolize L-ascorbate as a sole carbon source due to the reactivity of this molecule, and the generation of oxidative stress (7-12). Thus, aerobic catabolism of L-ascorbate requires the presence of specific amino acids (proline, threonine, or glutamine) in the culture medium that are proposed to decrease L-ascorbate oxidation (9, 10).

The gene cluster in *E. coli* responsible for L-ascorbate fermentation consists of the *ulaABCDEF* operon, and divergently transcribed genes *ulaG* and *ulaR* (7, 11, 13) (**Fig. 1A**). The *ulaABC* genes encode a phosphotransferase system (PTS) transporter for L-ascorbate uptake during both anaerobic and aerobic growth (9, 10, 12). The *ulaDEF* and *ulaG* genes are required for catabolism and *ulaR* encodes a regulator that acts as a repressor (7, 11, 13). L-ascorbate is converted to D-xylulose-5P, which can be further catabolized via the pentose phosphate pathway (7, 11, 13) (**Fig. 1B**). Regulation of the Ula system in *E. coli* is under the control of UlaR, cAMP-receptor protein (CRP), as well as integration host factor (IHF) (9, 10). Both IHF and UlaR act as repressors, while CRP is an activator of the *ulaABCDEF* operon (13). The Yia system shares three paralogous proteins with the Ula system, UlaD/YiaQ, UlaE/YiaR and UlaF/YiaS, which share less than 60% protein homology (**Fig. 1B**).

**Fig. 1.**
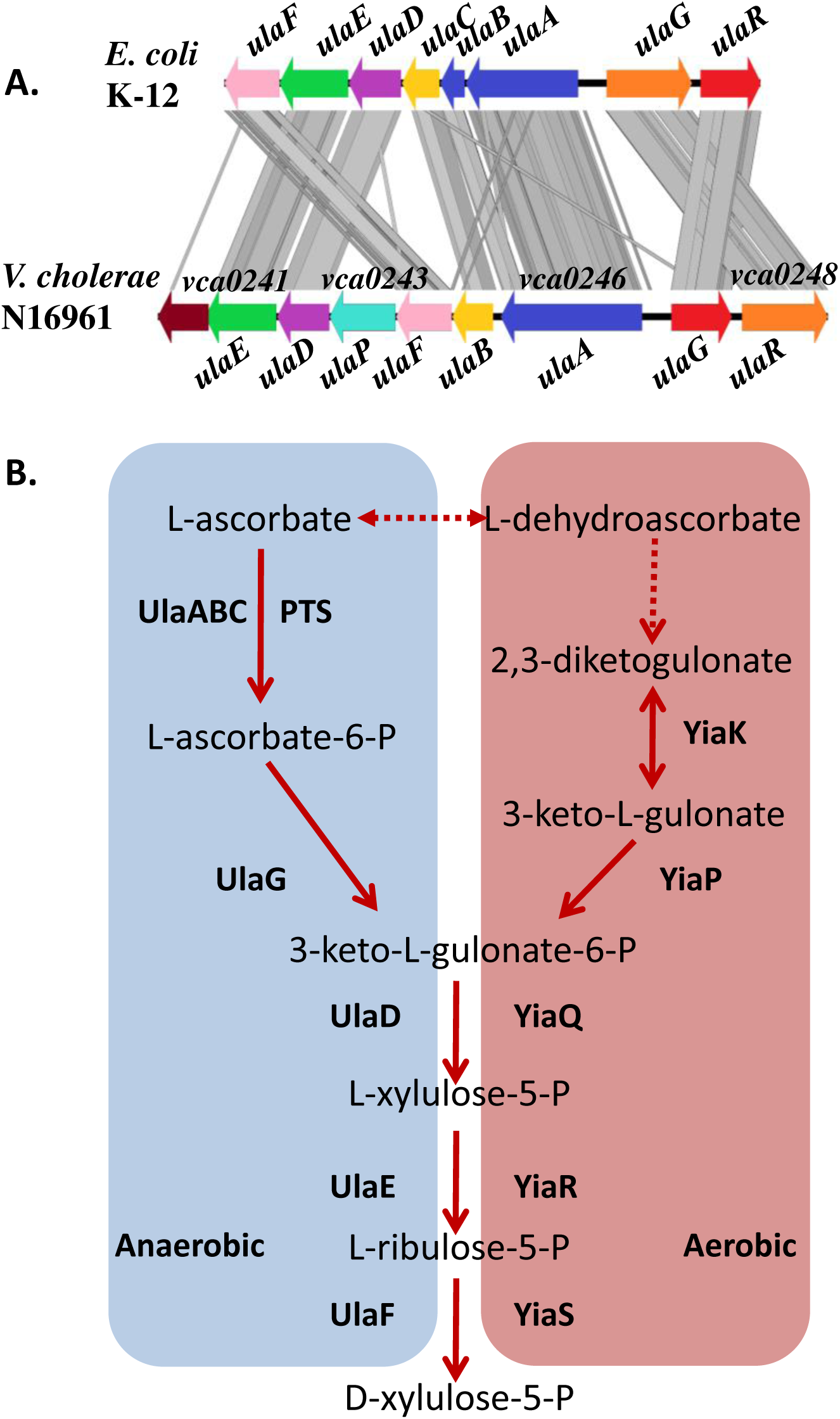
Pathways of L-ascorbate catabolism genes and pathways. **A**. Ula system gene order in *E. coli* K-12 and *V. cholerae* N16961. Gray bars indicate amino acid similarity via tBLASTx **B**. In anaerobic catabolism the transport and phosphorylation of L-ascorbate to L-ascorbate-6-phosphate occurs via *ulaABC*, which encodes a PTS-type transporter. L-ascorbate-6P is first converted to 3-keto-L-gulonate-6-P by the lactonase UlaG. Subsequent conversion to D-xylulose-5-P occurs via UlaDEF, which can be further catabolized by the Pentose Phosphate Pathway (PPP). In aerobic catabolism L-ascorbate is also transported into the cell via UlaABC. 2,3-diketogulonate is produced from L-ascorbate in the presence of oxygen and YiaK and YiaP can convert this to 3-keto-L-gulonate-6-P. YiaQRS are paralogs of UlaDEF.

*Vibrio cholerae* is a ubiquitous marine bacterium that is also an enteric pathogen of humans, and the causative agent of the deadly diarrheal disease, cholera. *V. cholerae* can colonize the small intestine of humans, where it produces cholera toxin that causes the secretory rice watery diarrhea, characteristic of cholera (14-17). Within the small intestine *V. cholerae* competes with the indigenous microbiota for nutrients and has to overcome many antagonistic interactions with the host microbiome (18-22). The nutrients *V. cholerae* utilize *in vivo* are poorly understood, but it was shown that this bacterium utilizes sialic acid, a component of intestinal mucus, as a sole carbon source and this ability has a significant fitness advantage *in vivo* (23-25). Additionally, *V. cholerae* was shown to use citrate fermentation *in vivo*, proposed to be an important phenotype for long-term carriage of *V. cholerae* (18).

In this study, we identified L-ascorbate fermentation *ula* gene clusters amongst *Vibrio* species, and demonstrated that *V. cholerae* can ferment L-ascorbate as a carbon and energy source. An in-frame deletion of *ulaG* (locus tag VCA0248) in *V. cholerae* was examined for growth on L-ascorbate. The expression pattern of *ula* genes in cells grown in the presence of L-ascorbate or intestinal mucus was determined. Additionally, we investigated whether L-ascorbate utilization resulted in a competitive advantage for cells grown in intestinal mucus. In *Aliivibrio fischeri, Vibrio* and *Photobacterium* species, within the *ula* operon, a gene (*vca0243*) that encodes a pyridoxal phosphate phosphatase homologous to YbhA from *Escherichia coli* is present. YbhA is an enzyme that converts pyridoxal phosphate (active vitamin C) to pyridoxal (inactive vitamin C). An in-frame deletion of *vca0243* in *V. cholerae* was examined to determine its effect on L-ascorbate catabolism and cellular metabolism in general. The role of CRP in L-ascorbate utilization was also investigated. Phylogenetic and genomic comparative analyses of UlaG, UlaD and the Ula cluster suggested that the region was acquired multiple times among *Vibrionaceae* and has a polyphyletic history.

## Results

### Ascorbate metabolism genes present in *Vibrio* species

The YiaQ protein from the YiaKLMNOPQRS cluster and UlaG from the Ula cluster of *E. coli* K-12 were used as seeds for BLASTp and psiBLAST analysis to identify L-ascorbate catabolism gene clusters in *V. cholerae* N16961. No proteins were identified with similarity to YiaQ or any other Yia protein within the *V. cholerae* N16961 genome. However, within chromosome II of *V. cholerae* N16961, we identified a homolog of UlaG (VCA0248), which is an L-ascorbic acid lactonase responsible for the first step in the catabolic pathway of L-ascorbate. Adjacent to *ulaG* was a *ulaR* homolog, which encoded a transcriptional repressor (7, 11, 13) (**Fig. 1A**). In *V. cholerae*, transcribed divergently from *ulaR-ulaG* was a set of seven ORFs, VCA0240-VCA0246, in a predicted operon, which shared homology with *ulaABCDEF* from *E. coli* (**Fig. 1A**). The first ORF VCA0246, was annotated as UlaA and was a fusion of the *ulaA* and *ulaB* genes from *E. coli*, which encoded components IIC and IIB of a phosphotransferase (PTS) system (**Fig. 1A**). VCA0245 shared homology with UlaC, the IIA component of the PTS system. VCA0242 and VCA0241 encoded homologs of UlaD and UlaE, respectively. In *E. coli*, UlaD is a putative decarboxylase that converts 3-keto-L-gulonate-6-P to L-xylulose-5P and UlaE is an epimerase that converts L-xylulose-5P to L-ribulose-5P (**Fig. 1B**) (7, 11). VCA0244 is a homolog of UlaF from *E. coli*, a 4-epimerase that converts _L_-ribulose-5P to D-xylulose-5-P (**Fig. 1B**). In *V. cholerae*, the Ula system contains an additional ORF, VCA0243 that is absent from the Ula system in enteric species. VCA0243 shares 40.44% percent identity, with 98% query cover and an E-value of 6e-62 to YbhA, a pyridoxal phosphate phosphatase in *E. coli*. A VCA0243 homolog was present within the Ula cluster in all *Vibrio* species examined.

### *Vibrio cholerae* and *V. vulnificus* utilization of L-ascorbate

Next, we determined whether L-ascorbate can be used as an alternative carbon and energy source by *Vibrio* species. To accomplish this, *V. cholerae* N16961, *V. vulnificus* CMCP6, *V. parahaemolyticus* RIMD2210633 and *E. coli* BW25113 were grown in microaerophilic conditions for 48 hours in M9 minimal media supplemented with casamino acids (M9CAS), M9CAS supplemented with 20mM glucose (M9CAS-Glucose), or M9CAS supplemented with 20mM L-ascorbate (M9CAS-Ascorbate). Casamino acids were added to growth media to alleviate the effects of L-ascorbate oxidation, which results in the production of reactive oxygen species (ROS) that are damaging to the cell (10). *Escherichia coli, V. cholerae* and *V. vulnificus* all grew significantly better in M9CAS-Ascorbate than M9CAS, indicating they can utilize L-ascorbate as an energy source (**Fig 2A-C**). As expected, *V. parahaemolyticus*, showed no difference between growth in M9CAS and M9CAS-Ascorbate as it does not contain the Ula gene cluster (**Fig. 2D**).

**Fig. 2.**
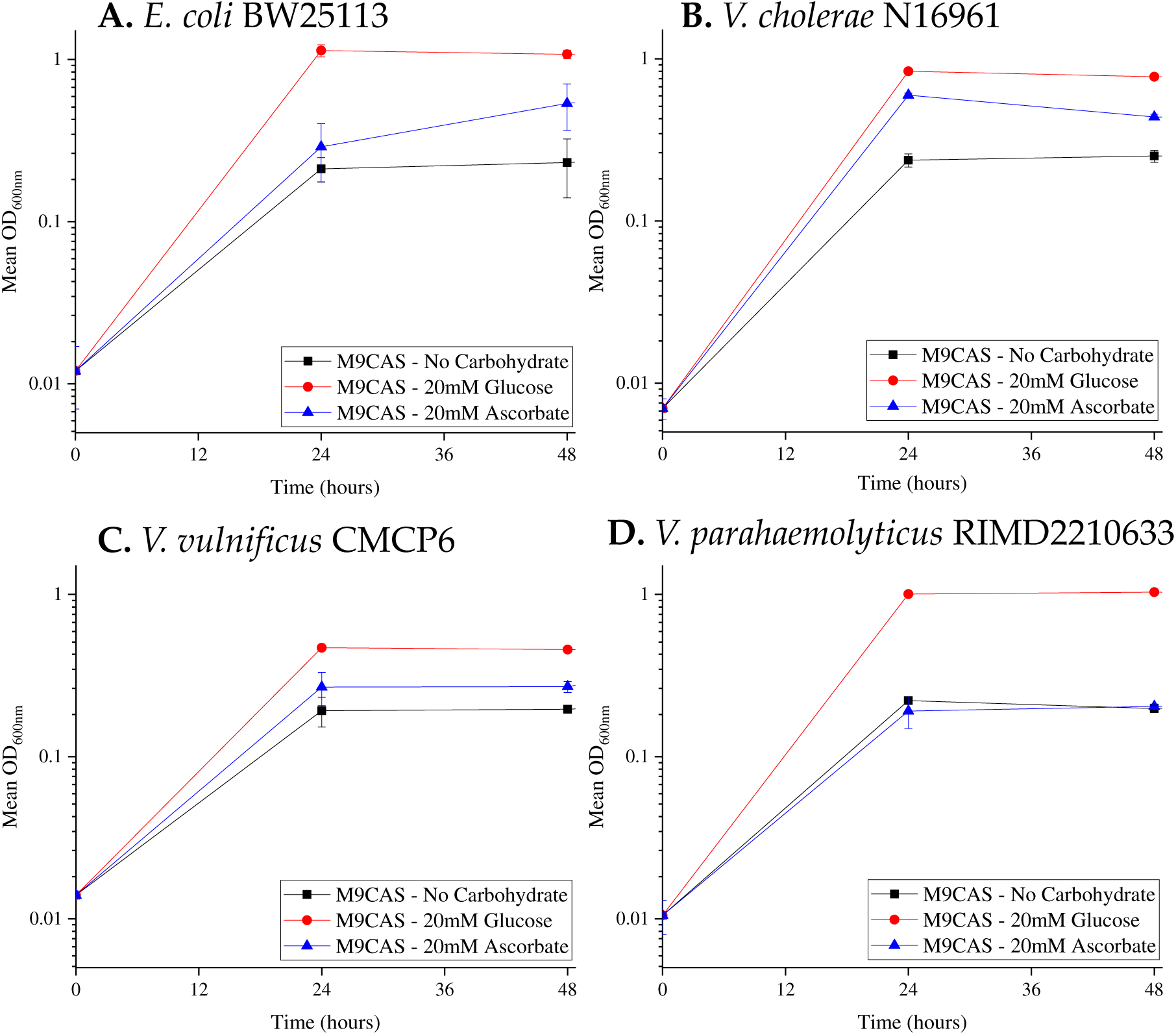
Growth analysis of *Vibrio* strains on L-ascorbate. (**A**) *E. coli* BW25113, (**B**) *V. cholerae* N16961, (**C**) *V. vulnificus* CMCP6 and (**D**) *V. parahaemolyticus* RIMD2210633 were grown in either M9CAS (squares), M9CAS-Glucose (circles), or M9CAS-ascorbate (triangles). Cells were grown statically under microaerophilic conditions for 48 hours. Error bars indicate standard deviation of the mean of two biological replicates.

### The *ulaG* gene is essential for L-ascorbate fermentation

We constructed an in-frame deletion of *ulaG* (VCA0247) in *V. cholerae* N16961 to investigate its requirement in L-ascorbate fermentation. The *ulaG* gene encodes an L-ascorbic acid lactonase, which is responsible for the first step in the L-ascorbate catabolic pathway. The wild type and Δ*ulaG* grown in LB 1%NaCl aerobically showed a similar pattern, indicating that the mutant had no overall growth defect (**Fig. S1**). Next, Δ*ulaG* and wild type were grown anaerobically for 48 hours in M9CAS, M9CAS-Glucose or M9CAS-Ascorbate. No growth defects were observed for Δ*ulaG* when grown in M9CAS or M9CAS-Glucose showing a similar growth pattern on both media (**Fig. S1**). However, Δ*ulaG* had a growth defect compared to wild type when grown in M9CAS-Ascorbate, with a significantly lower OD. These data show that *ulaG* is required for L-ascorbate fermentation in *V. cholerae* (**Fig. 3**).

**Fig. 3.**
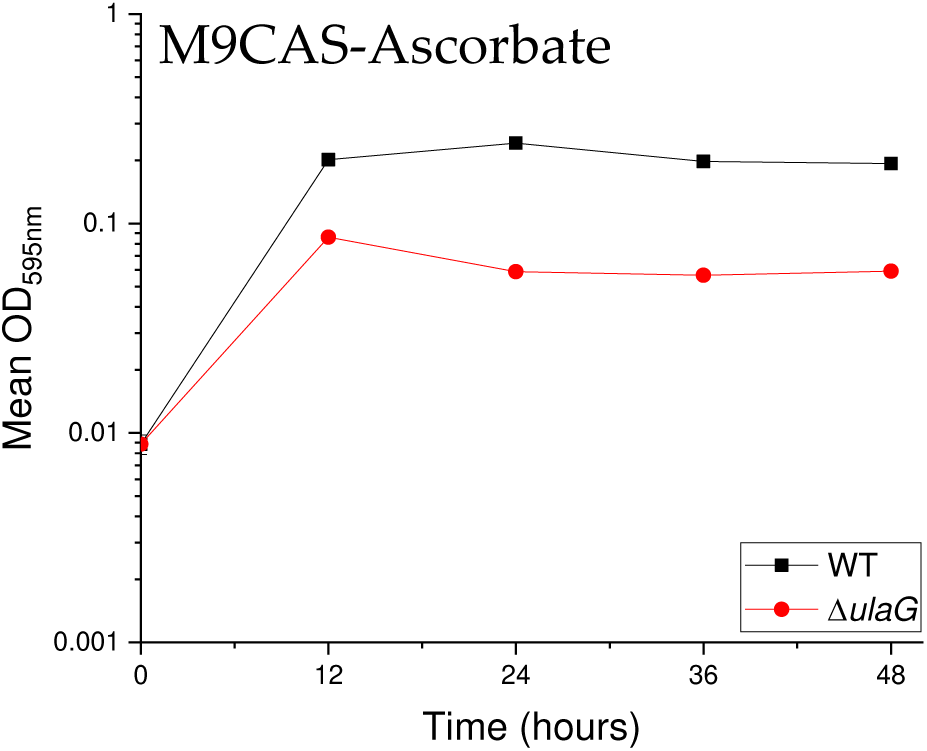
Growth analysis of *V. cholerae* N16961 and Δ*ulaG*. Wild type and Δ*ulaG* (Δ*vca0248*) were grown in M9CAS-Ascorbate statically under anaerobic conditions for 48 hours. Error bars indicate standard deviation of the mean of two biological replicates.

### The *ula* genes are induced in the presence of L-ascorbate and when grown on intestinal mucus

Expression of *ulaA* (*VCA0246*), *ulaR (VCA0247)* and *ulaG (VCA0248)* was analyzed in *V. cholerae* cells grown in M9CAS-Ascorbate, M9CAS-Glucose, M9CAS and M9-Mucus. In M9CAS-Ascorbate versus M9CAS-Glucose expression of *ulaA* was upregulated 788-fold, expression of *ulaR* was upregulated 568-fold, and *ulaG* was upregulated 1100-fold (**Fig. 4A**). To determine whether this induction was specific to L-ascorbate, and not due to catabolic repression by glucose, gene expression of cells grown in M9CAS-Ascorbate was also compared to cells grown in M9CAS without any additional carbohydrate supplementation. Under these conditions, the *ulaA, ulaR* and *ulaG* genes were significantly upregulated in M9CAS-Ascorbate compared to cells grown in M9CAS (**Fig. 4B**). Taken together, this data shows that the *ula* genes are highly induced in the presence of L-ascorbate.

**Fig. 4.**
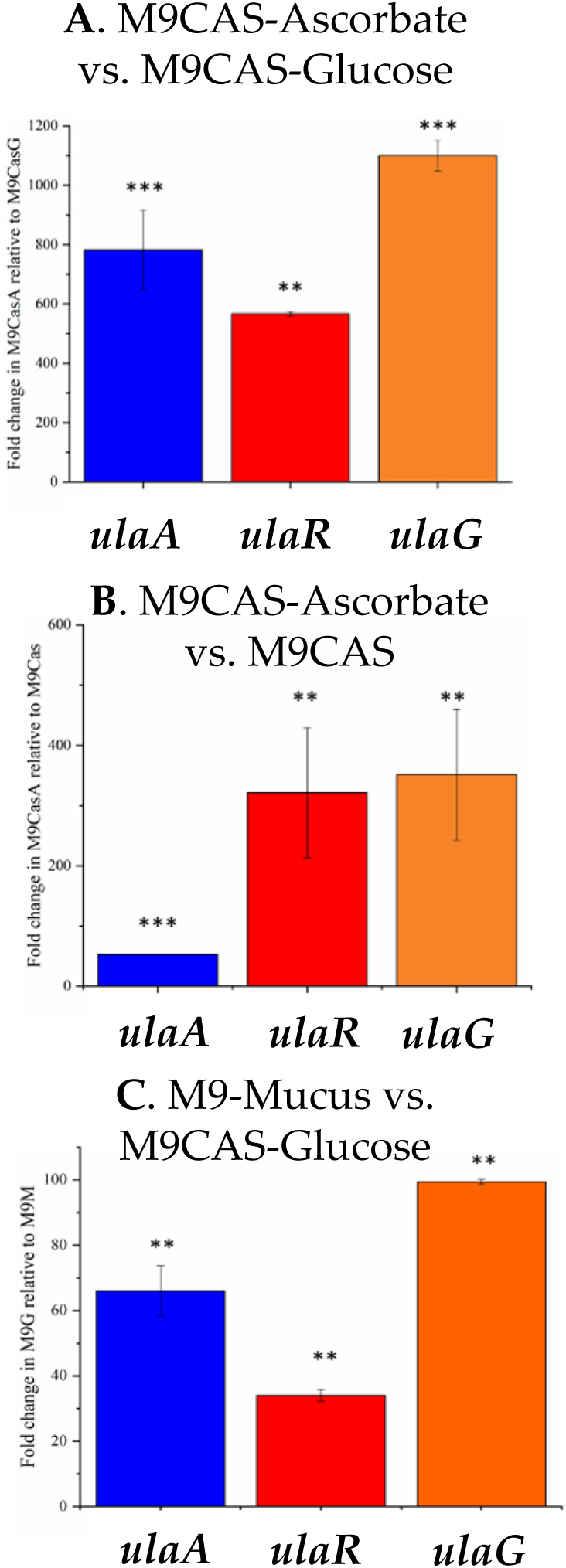
Expression analysis of L-ascorbate utilization genes. Expression analysis of *ulaA* (*vca0246*), *ulaR* (*vca0247*), and *ulaG* (*vca0248*) were performed from cells grown microaerophilically in **A**. M9CAS-Ascorbate relative to M9CAS-Glucose **B**. M9CAS-Ascorbate relative to M9CAS and **C**. M9-Mucus relative to cells M9CAS-Glucose. Data represent the mean fold change of two independent biological replicates. Error bars indicate standard deviation. Expression was normalized to the house keeping gene *topI*. A Student’s *t*-test was used to compare mean ΔC_T_ values between given conditions. * p<0.05, ** p<0.01, *** p<0.001.

Previous studies have shown that *Vibrio* species can utilize intestinal mucus as a carbon and energy source (25-28). Therefore, we examined the expression pattern of the *ula* genes in *V. cholerae* cells growth in M9-Mucus. The expression of *ulaA, ulaR*, and *ulaG* genes were significantly upregulated in cells grown in M9-mucus compared to M9CAS-glucose (**Fig. 4C**). Specifically, *ulaA* is upregulated 66-fold, *ulaR* is upregulated 34-fold and *ulaG* is upregulated 99-fold (**Fig. 4C**).

Next, we determined whether L-ascorbate fermentation provides a competitive advantage for *V. cholerae*. Fitness of the *ulaG* mutant relative to wild type was evaluated in *in vitro* competition assays in M9CAS-Ascorbate and M9CAS-mucus. Competitive indices (CI) for growth on M9CAS-Ascorbate was less than 0.1 indicating that the wild type outcompeted the *ulaG* mutant. Similarly, the CI in M9CAS-mucus was 0.55 demonstrating that the *ulaG* mutant was outcompeted by wild type suggesting that L-ascorbate fermentation may be an important *in vivo* phenotype (**Fig. 5**).

**Fig. 5.**
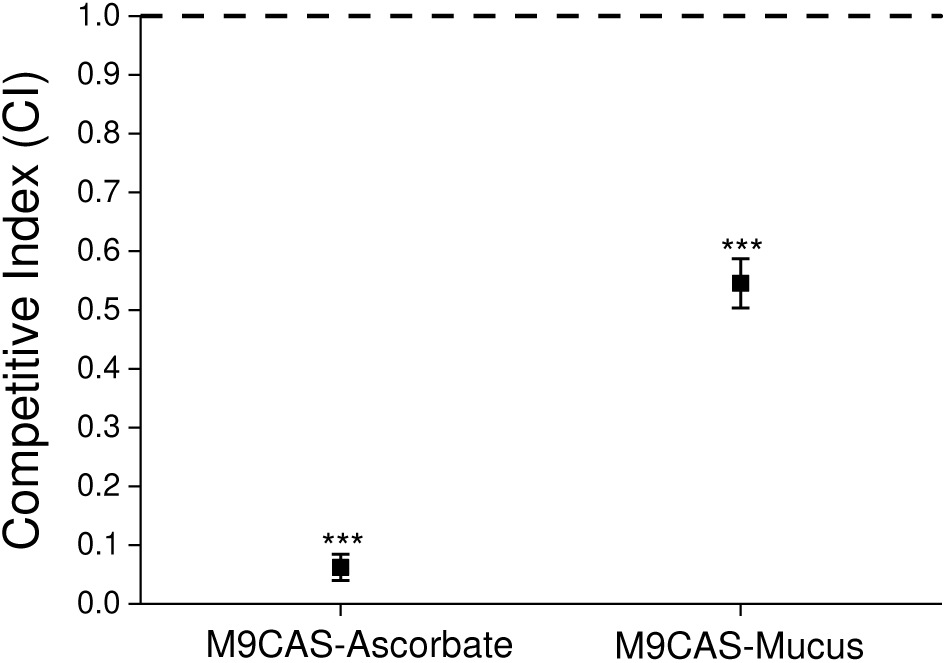
*In vitro* Competition assays. *V. cholerae ΔlacZ* and Δ*ulaG* strains were grown in 1:1 ratio in M9-Ascorbate or M9CAS-Mucus overnight and cells counted from a blue/white screen. Competitive Index (CI) was calculated using the formula: CI = output ratio _(Δ*ulaG*/ *ΔlacZ*)_ / input ratio _(Δ*ulaG* / *ΔlacZ*)_.

### VCA0243 a pyridoxal 5’ phosphate (PLP) phosphatase homolog

In *V. cholerae, vca0243* (AAF96154.1) encodes a 272 amino acid protein that shows 40.44% homology with 98% query cover to YbhA (VWQ01556.1) from *E. coli*. We constructed an in-frame deletion of *vca0243* in *V. cholerae* N16961 and examined growth in LB 1%NaCl. The mutant grew similar to wild type demonstrating no overall growth defect (**Fig. S2**). Next, we examined growth in M9CAS, M9CAS-Glucose and M9CAS-Ascorbate under anaerobic conditions. In M9CAS and M9CAS-glucose, the wild type and the mutant strains grew similar to one another (data not shown). In M9CAS-Ascorbate, the mutant had a growth defect compared to wild type with a significantly lower final OD (**Fig. 6A**) and in M9-Ascorbate, the mutant was unable to grow in the absence of CAS (**Fig. 6B**). These data indicates that *vca0243* is required for efficient L-ascorbate fermentation.

**Fig. 6.**
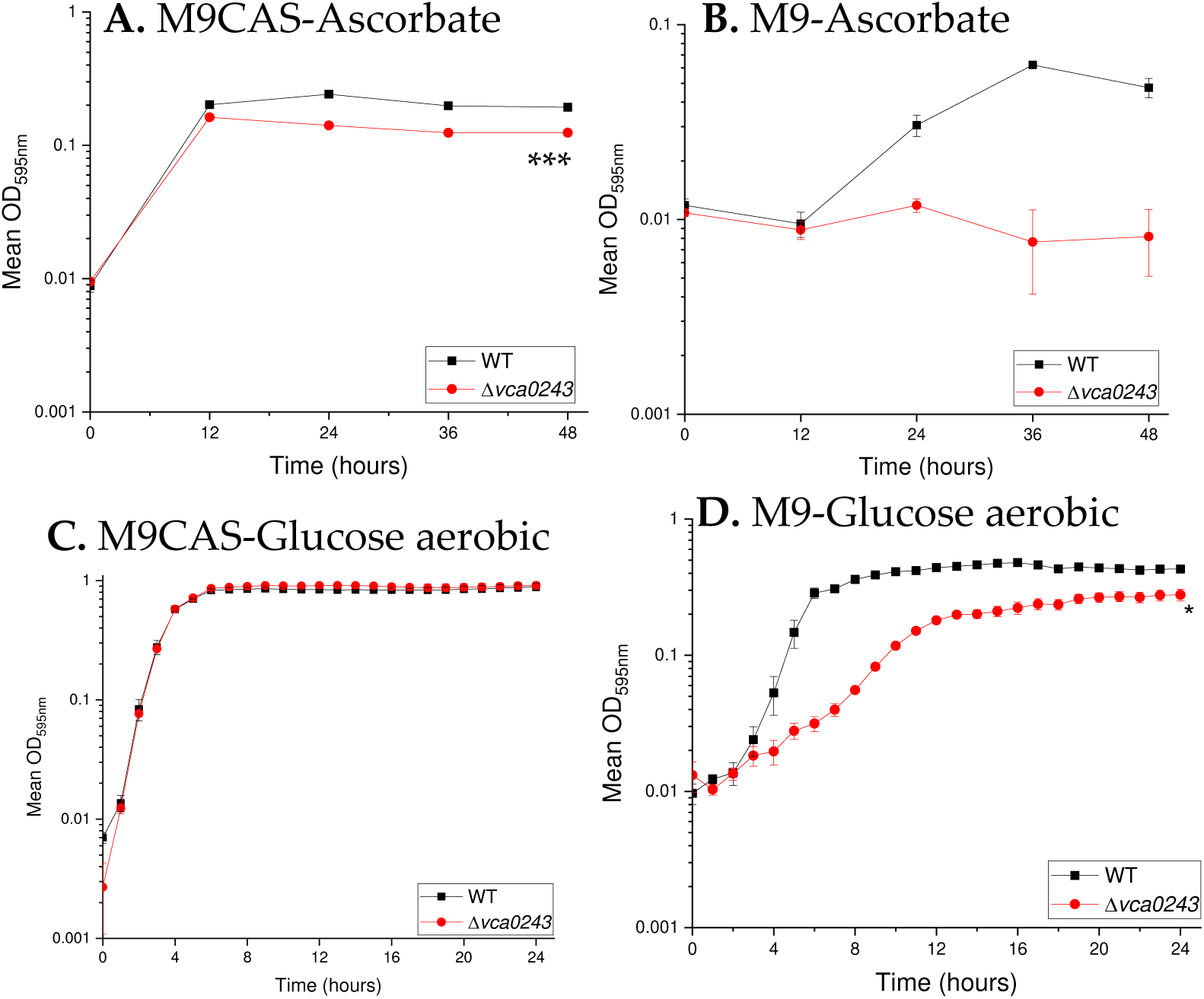
Growth analysis of *V. cholerae* N16961and Δ*vca0243*. Wild type and Δ*vca0243* were grown in **A**. M9CAS-Ascorbate or **B**. M9-Ascorbate under anaerobic conditions for 48 hours at 37°C. Wild type and Δ*vca0243* were grown in **C**. M9CAS-Glucose or **D**. M9-Glucose under aerobic conditions for 24 hours at 37°C. Error bars indicate standard deviation of the mean of two biological replicates. End-point biomass comparisons were performed using an unpaired Student’s *t*-test with a 95% confidence interval. *p<0.05 ***p<0.001

Deletion of *ybhA* in *E. coli* disrupted PLP homeostasis, causing growth inhibition and the presence of casamino acids was shown to alleviate this inhibition (29). Thus, we examined whether deletion of *vca0243* resulted in a growth defect in the absence of casamino acids when grown aerobically in glucose (**Fig. 6C and 6D**). In M9CAS-Glucose both wild type and the mutant grew similar, but in M9-Glucose, the mutant had a growth defect demonstrated by a longer lag phase and a reduced final OD (**Fig. 6D**). Overall, these data suggest that the VCA0243 is an required for optimal growth in the absence of casamino acids.

Next, we determined whether *vca0243* was important for growth on other carbon sources in general. To accomplish this, we used PM1 Biolog phenotype microarray growth assays to examine the utilization of 95 different carbon and energy sources by *V. cholerae* wild type and Δ*vca0243*. We found there were significant differences between wild type and Δ*vca0243* in 26 different carbon sources, which included amino acids or amino acid derivatives (**Fig. S3**). The area under the curve (AUC) of wild type growth was significantly higher than that of Δ*vca0243* in six of these amino acids, but significantly greater than wild type in L-threonine (**Fig. S3**). Differences in growth were also observed in 19 other carbon sources, with the wild-type strain utilizing these compounds significantly better than Δ*vca0243* (**Fig. S3**). Overall, the data indicate that VCA0243 has an important role in cellular metabolism.

### CRP is required for L-ascorbate fermentation in *V. cholerae*

It is known that cAMP-receptor protein (CRP) is a global regulator of alternative carbon metabolism in a variety of bacterial species (30-32). CRP was shown to be a regulator of the both the Yia and Ula system in *E. coli* and *K. pneumoniae* (9, 13). We constructed an in-frame deletion of *crp* (VC2614) and examined growth in M9CAS-Glucose. The *crp* mutant grew similar to wild type indicting no overall growth defect (**Fig. 7A**). Next, we examined growth inM9CAS-Ascorbate and show that Δ*crp* had a significant growth defect compared to wild type indicating that the mutant is unable to utilize L-ascorbate for growth (**Fig. 7B**).

**Fig. 7.**
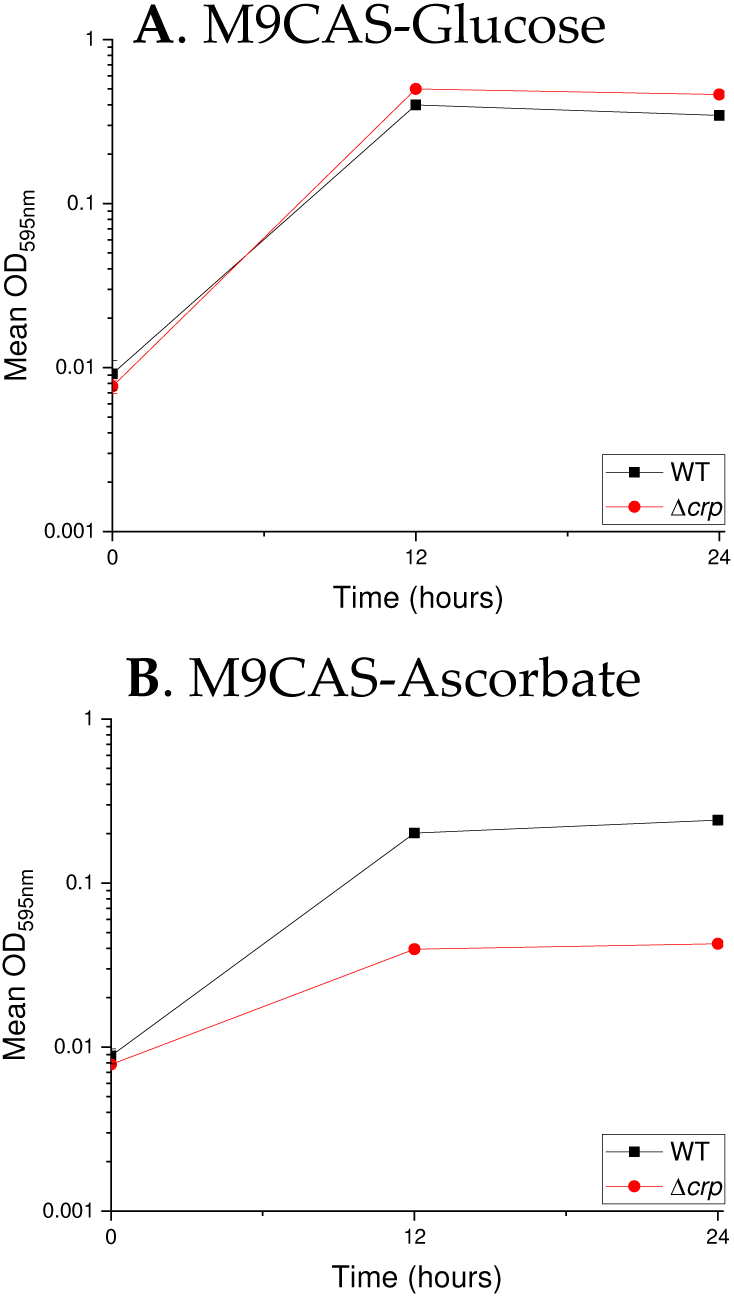
Growth analysis of *V. cholerae* N16961 and Δ*crp* on L-ascorbate. Wild type and Δ*crp* were grown in either **A**. M9CAS-Glucose or **B**. M9CAS-Ascorbic acid Cells were grown statically under anaerobic conditions for 24 hours. Error bars indicate standard deviation of the mean of two biological replicates.

**Fig. 8.** Evolutionary analysis of UlaG. The UlaG Maximum Likelihood tree with the highest log likelihood (−5835.54) is shown. The percentage of trees in which the associated taxa clustered together is shown next to the branches. Initial tree(s) for the heuristic search were obtained automatically by applying Neighbor-Join and BioNJ algorithms to a matrix of pairwise distances estimated using a JTT model, and then selecting the topology with superior log likelihood value. A discrete Gamma distribution was used to model evolutionary rate differences among sites (3 categories (+*G*, parameter = 0.3678)).

### Phylogenetic analysis of UlaG and UlaD among *Vibrionaceae*

To decipher the evolutionary history of ascorbate fermentation genes within the *Vibrionaceae*, we constructed a phylogenetic tree based on the UlaG and UlaD proteins. In addition, we constructed a tree based on the housekeeping protein RpoB from the same set of strains, as an indicator of the ancestral relationships among these species (33) (**Fig. S4**). On the UlaG tree, UlaG proteins that are present in members of the *Vibrionaceae* clustered within three major groups, which we designated A, B, and C, and encompassed 39 *Vibrio* species, *Aliivibrio fischeri* and 7 *Photobacterium* species (**Fig. 8**). Cluster A consists of UlaG from *V. cholerae* and its sister species, *V. mimicus and V. metoecus*. Also, within cluster A is UlaG from four members of the Campbellii clade, *V. harveyi, V. campbellii V. jasicida* and *V. owensii*, while UlaG is absent from all other members of this clade, which includes *V. parahaemolyticus* and *V. alginolyticus*.

Branching within cluster A, but divergently from *V. cholerae, V. mimicus* and *V. metoecus*, is UlaG from four *Providencia* species, members of the family *Morganellaceae*. However, the bootstrap value is low so this is likely not the origin of Ula from *Providencia*, but does suggest a close relationship with *Vibrio*. Of the four *Providencia* species that carry Ula, only *P. rettgeri* contained a *ybhA* homology within the *ula* cluster *ulaABybhAulaD*, and is present in all 30 *P. rettgeri* genomes in the database. In *P. alcalifaciens, P. rustigianii*, and *P. heimbachae*, the *ula* cluster consists of *ulaABD* with no *ybhA* homolog. In all four *Providencia* species the *ulaE* and *ulaF* homologs were present elsewhere on the genome (**Fig. S4**). The phylogenetic tree based on a housekeeping protein RpoB showed that the *Providencia* species clustered together divergent from *Vibrio* species, and are not closely related to *Vibrio* (**Fig S4**).

Within cluster B are *Vibrio* species belonging to the Splendidus clade, which cluster closely together; within this group is UlaG from *Aliivibrio fischeri* and *Photobacterium damselae* strains (**Fig. 8**). The positioning of UlaG from these species, nested within UlaG from *Vibrio* species, suggest the region has an origin in *Vibrio*. This is further evidenced by the RpoB tree showing that species from these genera cluster separately and distantly from *Vibrio* species (**Fig. S4**). Cluster C contained three groups of *Vibrio* species interspersed with UlaG from five *Photobacterium* species. The bootstrap values associated with *Photobacterium* species branching are extremely low indicating that the precise relationship with Vibrio species cannot be determined but does indicate a *Vibrio* origin for UlaG. Branching divergently from clusters A, B, and C is a single UlaG from *V. tritonius*.

The nearest relative to UlaG from *Vibrio* outside of cluster A, B and C is UlaG from *Yersinia ruckerii* and *Y. nurmii*, which branched together but divergently from *Vibrio* species. The next most closely related UlaG protein is from *Gilliamella apicola* and *G. apis*, gut symbionts of honeybees. *Gilliamella* is member of the *Orbaceae* family, a sister taxon to the order *Enterobacteriales* (34). In these species, the *ula* cluster is similar to *Vibrio* in terms of gene order but does not contain a *ybhA* homolog. The last group of UlaG proteins to cluster with UlaG from *Vibrio*, are UlaG from *Dickeya zeae, Yersinia intermedia* and five *Aeromonas* species. The *ula* genes are present in two divergently transcribed operons *ulaABFD* and *ulaGybhAER* in *Y. intermedia* and *Aeromonas* species, whereas in *D. zeae*, the *ula* genes are arranged *ulaABD* and *ulaGER* (**Fig. S4**). These are the only *Yersinia* and *Aeromonas* species, of the 12 described within *Yersinia* and the 14 within *Aeromonas* that contain the ascorbate fermentation gene cluster.

Given that *ulaG* and *ulaR* are divergently transcribed from the Ula transport and catabolism operon, we constructed a phylogenetic tree based on UlaD to determine whether the operon has a similar evolutionary history to UlaG. The UlaD tree showed some congruence among the genera in terms of their location within the tree (**Fig. S5**). However, the UlaD protein among *Vibrio* species showed low bootstrap values making comparisons among *Vibrio* species unreliable. UlaD from five *Photobacterium* species branched with the *Vibrio* UlaD proteins, but clustered at different locations with low bootstrap values. In addition, two *Photobacterium* species branched with UlaD from *Yersinia* and *Gilliamella* species but again the bootstrap value was very low indicating the exact placement of UlaD within the tree is hard to define. However, the placement of UlaD and UlaG proteins from *Photobacterium* species at different locations within the trees is in direct contrast to their branching together within the RpoB tree. *Aliivibrio fischeri* UlaD clustered with UlaD from the Splendidus clade, similar to UlaG and again this is in contrastto RpoB a locus ancestral to the species (**Fig. S4 and S5**). UlaD from *V. scophthalmi* was the most divergent protein examined in *Vibrio* and branched distantly from all other species. The UlaG protein from this species also branched divergently. Examination of the genome context of the Ula region in *V. scophthalmi* showed that the region was present on a plasmid pVS53 and thus was likely acquired by horizontal gene transfer.

Interestingly, analysis of L-ascorbate catabolism genes among members of the Splendidus clade identified a second YbhA homolog in these species (**Fig. S6**). This *ybhA* gene clustered with *yia* genes required for aerobic respiration of L-ascorbate and these genes were contiguous with the *ula* gene cluster (**Fig. S6**). This is a unique grouping of the aerobic and anaerobic L-ascorbate catabolism genes in these species and the presence of a YbhA homolog within each cluster suggests a critical role for YbhA in the L-ascorbate catabolism pathways.

## Discussion

In this study, we identified homologs of the Ula L-ascorbate fermentation cluster in *V. cholerae* and other *Vibrio* species. Growth pattern analysis revealed that *V. cholerae* and *V. vulnificus* could utilize L-ascorbate as an alternative nutrient source. In *V. cholerae*, we demonstrated that the *ulaG* gene is required for L-ascorbate fermentation and that the *ula* genes are induced in the presence of L-ascorbate. Our *in vitro* competition assays showed that Δ*ulaG* was outcompeted by wild type in M9CAS-Mucus indicting that L-ascorbate catabolism provides a fitness advantage. We also demonstrated that deletion of *crp* in *V. cholerae* resulted in a defect in growth, indicating that L-ascorbate fermentation is also under the control of CRP in this species.

Overall, out data suggest that L-ascorbate utilization is likely an important phenotype that may provide a fitness advantage *in vivo*. Previous studies in two human pathogens have suggested that ascorbate catabolism may be important for host colonization. Recent proteomics analysis of enterohemorrhagic *E. coli* (EHEC) in gnotobiotic piglets showed that some of the most differentially abundant proteins during infection were involved in L-ascorbate metabolism (35). Additionally, dual RNA-seq, RT-qPCR and metabolomics analysis of *H. ducreyi* in a human infection model demonstrated that genes associated with L-ascorbate recycling are upregulated in human hosts, while the most upregulated genes in *H. ducreyi* are the *ula* genes required for L-ascorbate fermentation (36).

We identified a homologue of a PLP phosphatase within the *ula* operon, encoded by VCA0243, that we showed is essential for growth on L-ascorbate in the absence of casamino acids, and also was required for efficient growth on several amino acids and sugars. PLP is a cofactor for many proteins and enzymes involved in a variety of cellular processes, including amino acid biosynthesis and degradation, as well as carbon metabolism (29, 37-39). It is widely known that PLP-dependent enzymes are associated with biochemical pathways that involve amino acids (39). A recent study in *E. coli* demonstrated that excess accumulation of either PL and PLP results in inhibition of growth initiation and changes in amino acid profiles (29). Growth initiation inhibition caused by PL or PLP accumulation was alleviated by addition of exogenous casamino acids to growth media (29). In *V. cholerae*, VCA0243 is likely a PLP phosphatase responsible for the dephosphorylation and inactivation of PLP, which could explain the growth defects of Δ*vca0243* in the absence of casamino acids. The remarkable clustering of both aerobic and anaerobic L-ascorbate catabolism genes in V. splendidus and related species and the presence of a YbhA homolog, which we named UlaP within each cluster, suggests the importance of PLP phosphatase in these pathways.

Our phylogenetic analysis showed that L-ascorbate fermentation genes are present in a subset of *Vibrio* species, a single *Aliivibrio* species, and seven *Photobacterium* species. For *Aliivibrio fischeri* and *Photobacterium* species, the UlaG and UlaD proteins clustered with *Vibrio* species suggests horizontal transfer among these groups.

Similarly, UlaG and UlaD proteins from *Providencia* species clustered with those from *Vibrio* species but the gene order is very different, indicating that this relationship is ancient. It is of interest to note that the *ula* genes from *A. fischeri* and *Providencia* species are located close to the *glmS* locus, which is the site of insertion of the transposon Tn7 in *E. coli*. In *Pr. heimbachae*, adjacent to the *ula* genes inserted at the *glmS* locus, is a Tn7 operon *tnsABCDE*. This may suggest a mode of acquisition of the *ula* genes in this species. In *Ph. damselae* strains that contain the *ula* cluster, the region is flanked on one side by a phage-like tyrosine recombinase integrase and the other end is flanked by phage genes and a restriction modification system. These genes are also present in *Ph. phosphoreum* strains that contain *ula* genes. In *Y. intermedia* directly adjacent to *ulaD* is a CRISPR repeat region and a type I-F CRISPR-Cas gene cluster followed by an integrase. In *A. veronii*, adjacent to *ulaD* is a tyrosine recombinase integrase. Finally, in *V. scophthalmi*, the *ula* genes were present on a plasmid again suggesting a mode of acquisition. Thus, along with our phylogenetic analysis, these signatures of horizontal gene transfer strongly suggest a polyphyletic origin for Ula and multiple acquisition events among species.

## Material and Methods

### Bacterial strains

*Vibrio cholerae* and *E. coli* were cultured aerobically with shaking (225rpm) in Luria Bertaini (LB) broth (Fisher Scientific, Waltham, MA) containing 1% NaCl at 37°C. *V. vulnificus* and *V. parahaemolyticus* were similarly cultured in LB containing a final concentration of 2% and 3% NaCl, respectively. Where indicated, M9 minimal media was utilized and supplemented with 0.02 mM MgSO_4_, and 0.1 mM CaCl_2_. Additionally, media was supplemented with 25 μg/mL of chloramphenicol (Cm) when required during mutant construction.

### Growth pattern analysis

*Vibrio cholerae* N16961 wild type and mutant strains were grown overnight at 37°C in LB 1% NaCl (LB 1%) with aeration, pelleted by centrifugation, washed with 1xPBS, and then resuspended in 1xPBS. Aerobic growth assays conducted in M9 minimal media containing 1% NaCl (M9) supplemented with 20mM glucose (M9-Glucose) or M9 containing 2% (w/v) casamino acids (M9CAS) (MP Biomedicals, LLC. Solon, OH) supplemented with 20mM glucose (M9CAS-Glucose) or with 20mM L-ascorbate (M9CAS-Ascorbate). Cultures were incubated at 37°C for 24 hours with intermittent shaking. OD_595nm_ was measured every hour utilizing a Tecan Sunrise (Tecan Group Ltd., Männedorf, Switzerland) microplate reader. A minimum of 2 biological replicates were performed for each condition. The mean OD_595nm_ and standard deviation of biological replicates was calculated for each assay. To create a microaerophilic environment, the cultures were aliquoted into fully filled snap-cap tubes and incubated statically at 37°C. At each time point, a sample was taken, centrifuged and resuspended in 1xPBS prior to measuring OD_600nm_. The concentration of NaCl was adjusted to 1% for *E. coli*, 2% for *V. cholerae* and *V. vulnificus*, and 3% for *V. parahaemolyticus* in each respective media. All microaerophilic growth assays included a minimum of three biological replicates. Anaerobic conditions were created by incubating microplates in airtight containers containing GasPak EZ Anaerobe sachets (BD Diagnostics, Sparks, MD). The OD_595nm_ of cultures was measured every 12 hours using a Tecan Sunrise spectrophotometric microplate reader. All assays were performed with a minimum of two biological replicates. The mean and standard deviation of biological replicates were calculated for each growth condition. End-point biomass comparisons were performed using an unpaired Students *t*-test with 95% confidence interval.

### Real time quantitative PCR (qPCR)

RNA was extracted from wild-type *V. cholerae* N16961 in M9 media supplemented with 0.02% (w/v) casamino acids (M9CAS), M9CAS+20 mM glucose (M9CAS-Glucose), or M9CAS+20 mM L-ascorbic acid (M9CAS-Ascorbate) and M9 supplemented with 20 mM glucose (M9-Glucose) or mouse intestinal mucus (M9-Mucus). Following 90 minutes of static growth at 37°C, RNA was isolated using TRIzol RNA extraction protocol according to the manufacturer’s instructions, and subsequently treated with Turbo DNase (Invitrogen). Purified RNA was quantified via Nanodrop spectrophotometer (Thermo Scientific, Waltham, MA) and 500 ng was used as a template for cDNA synthesis using Superscript III reverse transcriptase (Invitrogen). A 1:25 dilution of each cDNA sample was used for quantitative real-time PCR (qPCR). qPCR was carried out on a QuantStudio^™^ 6 Flex System with cycling parameters of a hold stage of 95°C for 20 s followed by a cycling stage of 95°C for 1 s and 60°C for 20 s. Relative gene expression was determined with Quantstudio Real-Time PCR Software v1.2 following the comparative C_T_ (ΔΔC_T_) method (40). The qPCR primers for *vca0246, vca0247*, and *vca0248* are listed in Table S1. Gene expression was normalized to the housekeeping gene for topoisomerase I (*topI*) listed in Table S1. RNA was isolated from two independent biological replicates and qPCR was performed in duplicate. Statistical analysis was performed comparing mean ΔC_T_ for a given target gene using a Student’s t-test with a 95% confidence interval

### In-frame deletion of *vca0243, ulaG* and *crp* (VC2614) construction

An in-frame *vca0243* deletion mutant was constructed via splicing by overlap-extension (SOE) and homologous recombination (41). The genome sequence of *V. cholerae* N16961 was used to design primers to create a 225-bp truncated *vca0243* PCR product. Gene fragments were amplified using primer pairs VCA0243_A/VCA0243_B and VCA0243_C/VCA0243_D, then digested via SacI and XbaI. The digested fragments were ligated with SacI and XbaI linearized pDS132, then transformed into *E. coli* DH5αλ*pir*.

Plasmid DNA was isolated from DH5α using the Nucleospin Plasmid kit (MACHEREY-NAGEL Inc., Bethlehem, PA) and transformed into the DAP-auxotroph donor strain *E. coli* β2155λ*pir*. The donor strain was conjugated overnight with the recipient strain *V. cholerae* N16961 on LB 1% plates. Selection of *V. cholerae* strains that had integrated deletion vectors into the genome was performed by plating conjugations on LB 1% plates containing Cm and confirmed by PCR. Colonies were then grown overnight without selection in LB 1% and plated on LB 1% supplemented with 10% sucrose (w/v) to select for strains that had undergone a second homologous recombination event. The genome sequence of *V. cholerae* N16961 was utilized to design primers for generation of a 110-bp truncated *ulaG* PCR product, and a 30-bp truncated *crp* PCR product. In-frame deletions of *ulaG* (*VCA0248*) and *crp* (*VC2614*) were constructed using a Gibson assembly protocol and HiFi DNA assembly master mix (New England Biolabs, Ipswich, MA). Gene fragments were amplified using primer pairs VC_ulaG_A/VC_ulaG_B and VC_ulaG_C/VC_ulaG_D or VC_CRP_A/VC_CRP_B and VC_CRP_C/VC_CRP_D. The resulting products were ligated and cloned into SacI linearized suicide-vector pDS132, then transformed into *E. coli* DH5α. Conjugation and selection of Δ*ulaG* and Δ*crp* mutants was performed in the same manner as described for the Δ*vca0243* mutant. Primers used for construction of deletion mutants are listed in Table S1. Restriction enzyme cut sites of SOE PCR primers are indicated by lowercase, italicized letters. Complementary regions for Gibson assembly and SOE PCR are represented by lowercase letters in each primer sequence. All primers utilized in this study were purchased from IDT (Integrated DNA Technologies, Coralville, IA). Deletion mutants were confirmed via PCR and DNA sequencing.

### *In vitro* competition assays

A *V. cholerae* Δ*lacZ* strain was utilized as a surrogate for wild-type, as deletion of *lacZ* allows for subsequent blue/white colony selection when plated on X-gal LB 1% plates. This mutant was shown to have no other defect compared to wild type (23). *V. cholerae* Δ*lacZ* and genetic mutants were grown aerobically in LB 1% for 24 hours, pelleted by centrifugation, washed and resuspended in PBS at a concentration of 1⨯10^10^ CFU/mL. Cultures were then diluted in a 1:1 ratio (mutant:Δ*lacZ*) and inoculated 1:50 into fully-filled and parafilmed 15mL conical tubes containing M9CAS-Ascorbate or M9-CAS supplemented with 30 μg/mL mouse intestinal mucus. Cultures were incubated anaerobically at 37^°^C in airtight containers with GasPak EZ anaerobe sachets for 24 hours, then serially diluted and plated on X-gal LB 1% plates. The competitive index (CI) of each strain was calculated using the formula: CI = output ratio _(Mutant / N16961Δ*lacZ*)_ / input ratio _(Mutant / N16961Δ*lacZ*)_. The ratio of mutant strain to Δ*lacZ* in the inoculum was referred to as the “input ratio”, while the ratio of mutant colonies to Δ*lacZ* colonies observed after competition was referred to as the “output ratio”. If an assay resulted in a CI > 1.0, the mutant outcompeted the wild-type; while a CI < 1.0 indicated that the wild-type strain outcompeted the mutant.

### Bioinformatics analysis

The YiaS and UlaG proteins of *E. coli* K-12 (NP_418613.2) were used as seeds for BLASTp analysis against *V. cholerae* to identify putative L-ascorbate catabolism proteins. The identified *V. cholerae* UlaG homolog VCA0248 (AWA79938.1) was subsequently seeded for BLASTp analysis against other *Vibrionaceae* species.

Additionally, BLASTp analysis of the UlaA homolog VCA0246 (NP_232644.1) from *V. cholerae* was performed against *Vibrionaceae* representative species in the NCBI genome database. Analysis of VCA0243 (AAF96154.1, AWB75675.1) was also conducted using BLASTp against *Vibrionaceae* and non-*Vibrio* species. Identified genomic regions were downloaded from the National Center for Biotechnology Information (NCBI) database. Figures representing amino acid similarity and gene cluster arrangement were generated from genome sequences using tBLASTx in Easyfig 2.2.3 (42). Structural analysis of VCA0243 by HHpred (43) was conducted using the protein FASTA sequence (AAF96154.1) from NCBI.

### Growth phenotype assays

Growth characteristics of *V. cholerae* N16961 and Δ*vca0243* strains were examined on various carbon sources via a PM1 phenotype microarray (Biolog, Inc., Hayward, CA). Briefly, aerobically grown LB 1% overnight cultures were pelleted by centrifugation, washed with 1xPBS and resuspended in M9 minimal containing 1% NaCl (M9). Cultures were then diluted 1:100 in M9, and 100μL was inoculated into each well of a PM1 Biolog phenotype microarray. Cultures were incubated at 37^°^C for 24 hours with intermittent shaking and the OD_595nm_ was measured every hour utilizing a Tecan Sunrise microplate reader. Following assay completion, the area under the curve (AUC) of each strain was calculated using Origin 2019 (OriginLab Corporation, Northampton, MA). The AUC of the negative control (blank) was subtracted from the AUC in each respective media, and the mean AUC of wild-type and Δ*vca0243* were compared using an unpaired Student’s *t*-test with 95% confidence interval. A minimum of two biological replicates was performed for each strain.

### Phylogenetic analyses

Phylogenetic analysis was conducted using UlaG (VCA0246) and UlaD (VCA0240) protein as a seed to identity all homologs within the family *Vibrionaceae* and their closest relatives. Unique protein sequences that had >95% sequence coverage and >70% amino acid identity with UlaG and UalD were obtained from NCBI database and aligned using the ClustalW (44). The housekeeping protein RpoB was used to show the evolutionary relationships among all the species examined in the UlaG and UlaD trees. The evolutionary history of each protein was inferred by using the Maximum Likelihood method and Le_Gascuel_2008 model as determine by best fit model selection in MEGAX (45, 46). The trees are drawn to scale, with branch lengths measured in the number of substitutions per site. UlaG analysis involved 68, UlaD 63 and RpoB 64 amino acid sequences. All positions with less than 95% site coverage were eliminated, i.e., fewer than 5% alignment gaps, missing data, and ambiguous bases were allowed at any position (partial deletion option). There were a total of 354 positions for UlaG, 214 positions for UlaD, 1341 positions for RpoB in the final dataset. Evolutionary analyses were conducted in MEGA (46).

## ACKNOWLEDGEMENTS

This research was supported by a National Science Foundation grant (award IOS-1656688) to E.F.B. We thank members of the Boyd Group for constructive feedback on the manuscript.

---

**Table 1.**
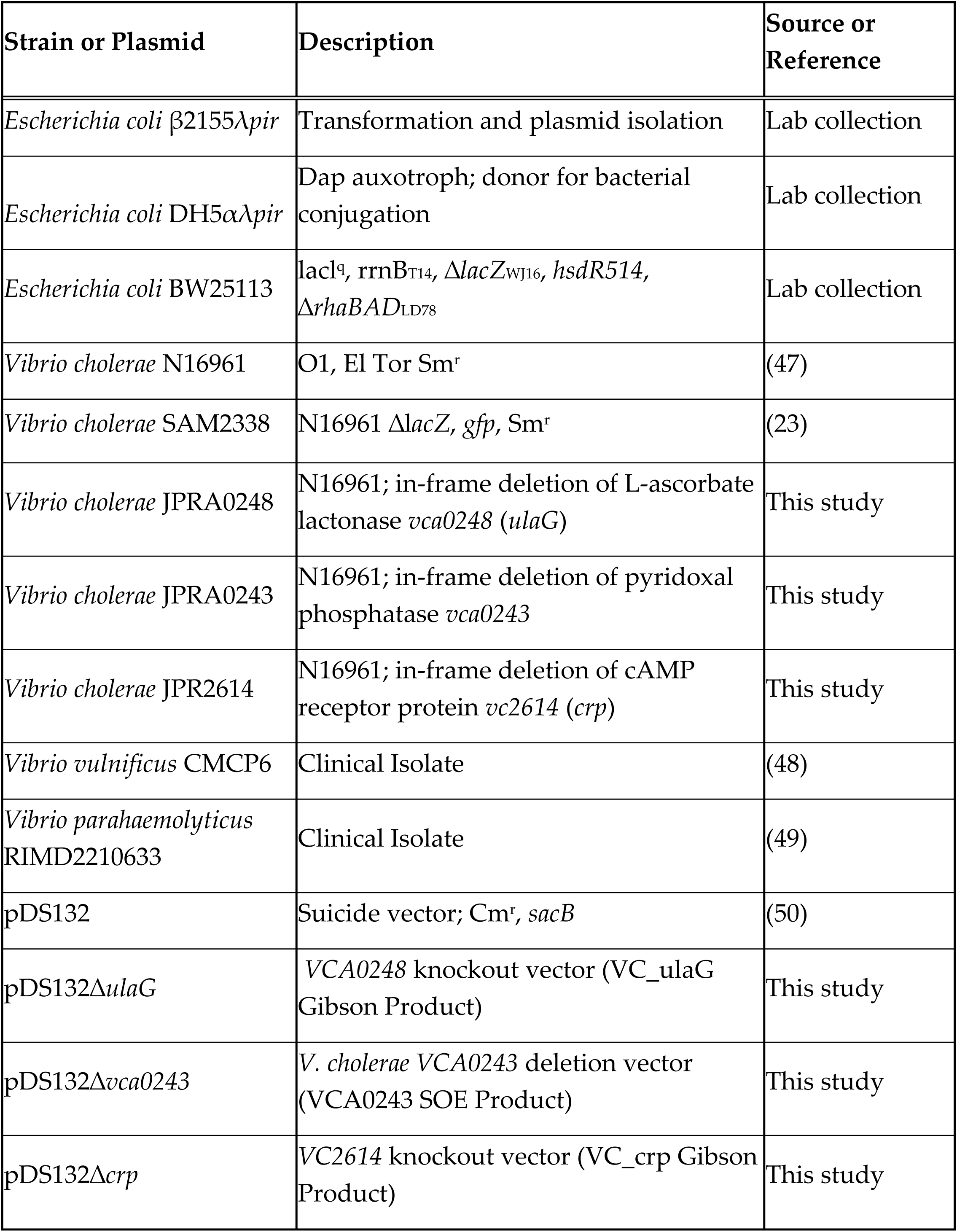
Bacterial strain and plasmids used in this study

**Table 2.**
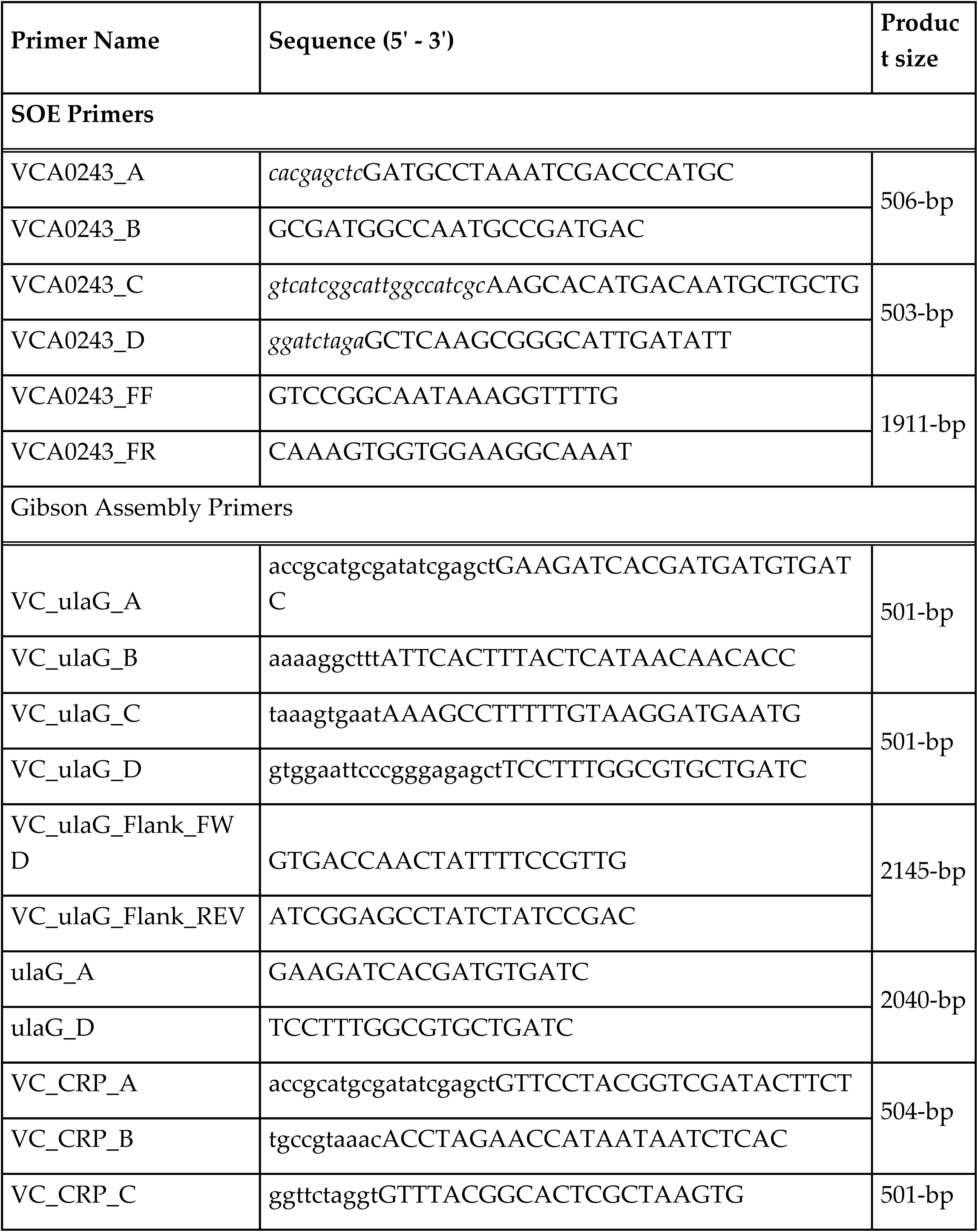
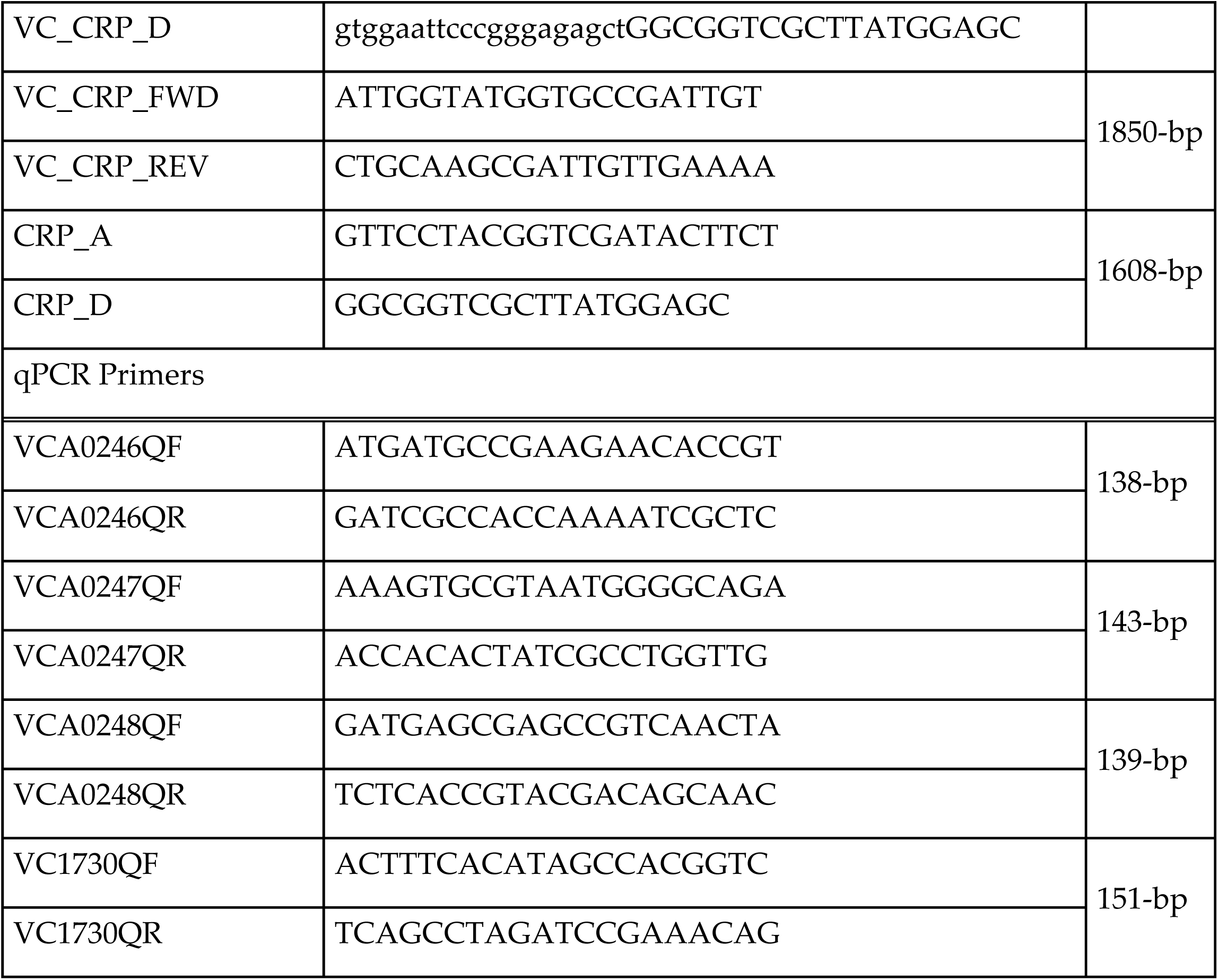
Primers

## Supplemental Data

**Fig. S1. A. Growth of *V. cholerae* wild type and Δ*ulaG*. A**. Growth in LB 1%NaCl. for 24 hours. Growth in B. M9CAS or C. M9CAS-Glucose anaerobically for 48 hours. Error bars indicate standard deviation of the mean of two biological replicates.

**Fig. S2. Growth of *V. cholerae* wild type and Δ*vca0243*** in LB aerobically Cells were grown 24 hours. Error bars indicate standard deviation of the mean of two biological replicates.

**Fig. S3. Growth of *V. cholerae* N16961 and Δ*vca0243* on various carbon sources**. *V. cholerae* N16961 (black bars) and Δ*vca0243* (white bars) were diluted in M9 minimal media and grown aerobically for 24 hours on various carbon sources in a PM1 Biolog Phenotype Microarray. Carbon utilization results of non-amino acid sources are included in **A**-**C**. Use of amino acids and derivatives are displayed in **D**. Mean AUC was calculated for each carbon source in Origin 2019. Error bars indicate standard error of the mean of two biological replicates. Mean AUC comparison was performed utilizing an unpaired Students *t*-test with 95% confidence interval. *p<0.05, **p<0.01, ***p<0.001. (N-A-G: *N*-acetylglucosamine).

**Fig. S4. A. *Vibrionaceae* housekeeping tree based on RpoB using the Neighbor-Joining method**. The optimal tree with the sum of branch length = 1.18 is shown. The bootstrap test (500 replicates) values are shown next to the branches. The evolutionary distances were computed using the JTT matrix-based method and are in the units of the number of amino acid substitutions per site. The rate variation among sites was modeled with a gamma distribution (shape parameter = 5). This analysis involved 64 amino acid sequences. There were a total of 1341 positions in the final dataset. Bacterial families to which each species belongs are labelled and major clades of *Vibrio* species are named in orange. **B**. The *ula* gene clusters present among the strains examined. Arrows indicate ORFs and the direction of transcription and black lines indicate genes are contiguous. Homologous genes are colored similarly.

**Fig. S5. Evolutionary analysis of UlaD among Vibrionaceae**. UlaD Neighbor-joining tree. The optimal tree with the sum of branch length = 3.8 is shown. Bootstrap test (500 replicates) values are shown next to the branches. The evolutionary distances were computed using the JTT matrix-based method and are in the units of the number of amino acid substitutions per site. The rate variation among sites was modeled with a gamma distribution (shape parameter = 5). This analysis involved 63 amino acid sequences. There were a total of 214 positions in the final dataset.

**Fig. S6. Unique clustering of aerobic and anaerobic L-ascorbate catabolism operons**. The Yia/Ula gene clusters group together on the genomes of species from the Spendidus clade. Arrow depict ORFs and direction of arrows indicate direction of transcription. Purple arrows labelled YbhA indicate the two copies of the pyroxidal phosphatase.

---

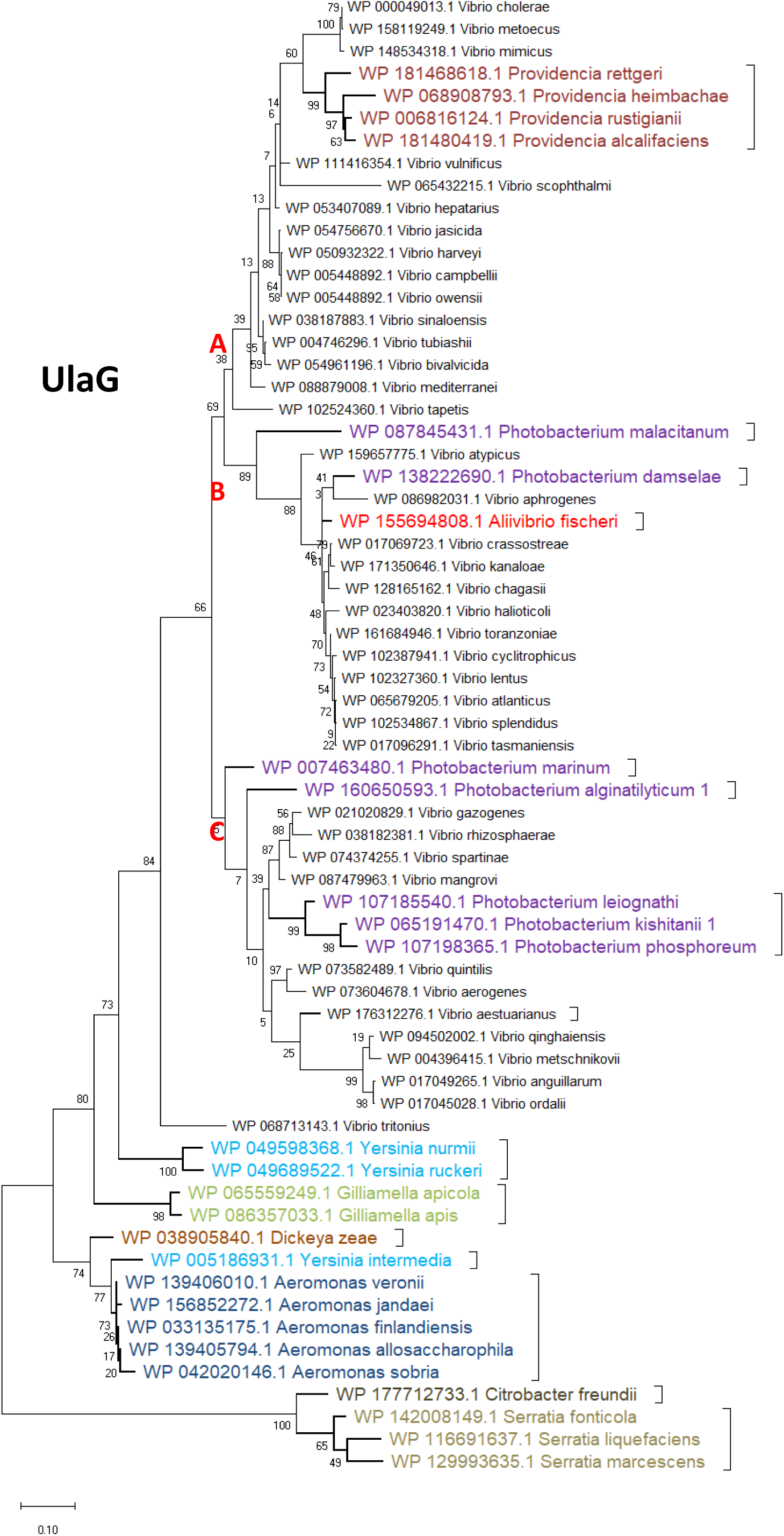

## References

1. Traber M, Stevens J. 2011. Vitamins C and E: beneficial effects from a mechanistic perspective. Free Radic Biol Med 51:1000–13..

2. Granger M, Eck P. 2018. Dietary Vitamin C in Human Health. Adv Food Nutr Res 83:281–310.

3. Carr AC, Maggini S. 2017. Vitamin C and Immune Function. Nutrients 9.

4. Chatterjee IB, Kar NC, Ghosh NC, Guha BC. 1961. Aspects of ascorbic acid biosynthesis in animals. Ann N Y Acad Sci 92:36–56.

5. Drouin G, Godin JR, Pagé B. 2011. The genetics of vitamin C loss in vertebrates. Curr Genomics 12:371–8.

6. Henriques SF, Duque P, Lopez-Fernandez H, Vazquez N, Fdez-Riverola F, Reboiro-Jato M, Vieira CP, Vieira J. 2019. Multiple independent L-gulonolactone oxidase (GULO) gene losses and vitamin C synthesis reacquisition events in non-Deuterostomian animal species. BMC Evol Biol 19:126.

7. Campos E, Aguilar J, Baldoma L, Badia J. 2002. The gene yjfQ encodes the repressor of the yjfR-X regulon (ula), which is involved in L-ascorbate metabolism in Escherichia coli. J Bacteriol 184:6065–8.

8. Campos E, Aguilera L, Gimenez R, Aguilar J, Baldoma L, Badia J. 2009. Role of YiaX2 in L-ascorbate transport in Klebsiella pneumoniae 13882. Can J Microbiol 55:1319–22.

9. Campos E, de la Riva L, Garces F, Gimenez R, Aguilar J, Baldoma L, Badia J. 2008. The yiaKLX1×2PQRS and ulaABCDEFG gene systems are required for the aerobic utilization of L-ascorbate in Klebsiella pneumoniae strain 13882 with L-ascorbate-6-phosphate as the inducer. J Bacteriol 190:6615–24.

10. Campos E, Montella C, Garces F, Baldoma L, Aguilar J, Badia J. 2007. Aerobic L-ascorbate metabolism and associated oxidative stress in Escherichia coli. Microbiology (Reading) 153:3399–3408.

11. Yew WS, Gerlt JA. 2002. Utilization of L-ascorbate by Escherichia coli K-12: assignments of functions to products of the yjf-sga and yia-sgb operons. J Bacteriol 184:302–6.

12. Zhang Z, Aboulwafa M, Smith MH, Saier MH, Jr. 2003. The ascorbate transporter of Escherichia coli. J Bacteriol 185:2243–50.

13. Campos E, Baldoma L, Aguilar J, Badia J. 2004. Regulation of expression of the divergent ulaG and ulaABCDEF operons involved in LaAscorbate dissimilation in Escherichia coli. J Bacteriol 186:1720–8.

14. Colwell RR, Huq A, Chowdhury MA, Brayton PR, Xu B. 1995. Serogroup conversion of *Vibrio cholerae*. Can J Microbiol 41:946–50.

15. Faruque SM, Albert MJ, Mekalanos JJ. 1998. Epidemiology, genetics, and ecology of toxigenic Vibrio cholerae. Microbiol Mol Biol Rev 62:1301–14.

16. Thelin K, Taylor R. 1996. Toxin-coregulated pilus, but not mannose-sensitive hemagglutinin, is required for colonization by *Vibrio cholerae* O1 El Tor biotype and O139 strains. Infect Immun 64:2853–2856..

17. Waldor MK, Mekalanos JJ. 1996. Lysogenic conversion by a filamentous phage encoding cholera toxin. Science 272:1910–4.

18. Liu M, Hao G, Li Z, Zhou Y, Garcia-Sillas R, Li J, Wang H, Kan B, Zhu J. 2019. CitAB Two-Component System-Regulated Citrate Utilization Contributes to Vibrio cholerae Competitiveness with the Gut Microbiota. Infect Immun 87.

19. Wang J, Xing X, Yang X, Jung IJ, Hao G, Chen Y, Liu M, Wang H, Zhu J. 2018. Gluconeogenic growth of Vibrio cholerae is important for competing with host gut microbiota. J Med Microbiol 67:1628–1637.

20. Midani FS, Weil AA, Chowdhury F, Begum YA, Khan AI, Debela MD, Durand HK, Reese AT, Nimmagadda SN, Silverman JD, Ellis CN, Ryan ET, Calderwood SB, Harris JB, Qadri F, David LA, LaRocque RC. 2018. Human Gut Microbiota Predicts Susceptibility to Vibrio cholerae Infection. J Infect Dis 218:645–653.

21. Logan SL, Thomas J, Yan J, Baker RP, Shields DS, Xavier JB, Hammer BK, Parthasarathy R. 2018. The Vibrio cholerae type VI secretion system can modulate host intestinal mechanics to displace gut bacterial symbionts. Proc Natl Acad Sci U S A 115:E3779–E3787.

22. Zhao W, Caro F, Robins W, Mekalanos JJ. 2018. Antagonism toward the intestinal microbiota and its effect on Vibrio cholerae virulence. Science 359:210–213.

23. Almagro-Moreno S, Boyd EF. 2009. Sialic acid catabolism confers a competitive advantage to pathogenic vibrio cholerae in the mouse intestine. Infect Immun 77:3807–16.

24. Almagro-Moreno S, Boyd EF. 2009. Insights into the evolution of sialic acid catabolism among bacteria. BMC Evol Biol 9:118.

25. McDonald ND, Lubin JB, Chowdhury N, Boyd EF. 2016. Host-Derived Sialic Acids Are an Important Nutrient Source Required for Optimal Bacterial Fitness In Vivo. mBio 7:e02237–15.

26. Reddi G, Pruss K, Cottingham KL, Taylor RK, Almagro-Moreno S. 2018. Catabolism of mucus components influences motility of Vibrio cholerae in the presence of environmental reservoirs. PLoS One 13:e0201383.

27. Kalburge SS, Carpenter MR, Rozovsky S, Boyd EF. 2017. Quorum Sensing Regulators Are Required for Metabolic Fitness in Vibrio parahaemolyticus. Infect Immun 85.

28. Whitaker WB, Richards GP, Boyd EF. 2014. Loss of sigma factor RpoN increases intestinal colonization of Vibrio parahaemolyticus in an adult mouse model. Infect Immun 82:544–56.

29. Sugimoto R, Saito N, Shimada T, Tanaka K. 2018. Identification of YbhA as the pyridoxal 5’-phosphate (PLP) phosphatase in Escherichia coli: Importance of PLP homeostasis on the bacterial growth. J Gen Appl Microbiol 63:362–368.

30. Savery N, Rhodius V, Busby S. 1996. Protein-protein interactions during transcription activation: the case of the Escherichia coli cyclic AMP receptor protein. Philos Trans R Soc Lond B Biol Sci 351:543–50.

31. Colton DM, Stabb EV. 2016. Rethinking the roles of CRP, cAMP, and sugar-mediated global regulation in the Vibrionaceae. Curr Genet 62:39–45.

32. Green J, Stapleton MR, Smith LJ, Artymiuk PJ, Kahramanoglou C, Hunt DM, Buxton RS. 2014. Cyclic-AMP and bacterial cyclic-AMP receptor proteins revisited: adaptation for different ecological niches. Curr Opin Microbiol 18:1–7.

33. Boyd EF, Carpenter MR, Chowdhury N, Cohen AL, Haines-Menges BL, Kalburge SS, Kingston JJ, Lubin JB, Ongagna-Yhombi SY, Whitaker WB. 2015. Post-Genomic Analysis of Members of the Family Vibrionaceae. Microbiol Spectr 3.

34. Kwong WK, Moran NA. 2013. Cultivation and characterization of the gut symbionts of honey bees and bumble bees: description of Snodgrassella alvi gen. nov., sp. nov., a member of the family Neisseriaceae of the Betaproteobacteria, and Gilliamella apicola gen. nov., sp. nov., a member of Orbaceae fam. nov., Orbales ord. nov., a sister taxon to the order ‘Enterobacteriales’ of the Gammaproteobacteria. Int J Syst Evol Microbiol 63:2008–2018.

35. Pieper R, Zhang Q, Clark DJ, Parmar PP, Alami H, Suh MJ, Kuntumalla S, Braisted JC, Huang ST, Tzipori S. 2013. Proteomic View of Interactions of Shiga Toxin-Producing Escherichia coli with the Intestinal Environment in Gnotobiotic Piglets. PLoS One 8:e66462.

36. Griesenauer B, Tran TM, Fortney KR, Janowicz DM, Johnson P, Gao H, Barnes S, Wilson LS, Liu Y, Spinola SM. 2019. Determination of an Interaction Network between an Extracellular Bacterial Pathogen and the Human Host. mBio 10.

37. Richts B, Rosenberg J, Commichau FM. 2019. A Survey of Pyridoxal 5’-Phosphate-Dependent Proteins in the Gram-Positive Model Bacterium Bacillus subtilis. Front Mol Biosci 6:32.

38. Percudani R, Peracchi A. 2003. A genomic overview of pyridoxal-phosphate-dependent enzymes. EMBO Rep 4:850–4.

39. Schneider G, Käck H, Lindqvist Y. 2000. The manifold of vitamin B6 dependent enzymes. Structure 8:R1–6.

40. Pfaffl MW. 2001. A new mathematical model for relative quantification in real-time RT-PCR.. Nucleic Acids Res 29:e45.

41. Horton RM, Hunt HD, Ho SN, Pullen JK, Pease LR. 1989. Engineering hybrid genes without the use of restriction enzymes: gene splicing by overlap extension. Gene 77:61–8.

42. Sullivan MJ, Petty NK, Beatson SA. 2011. Easyfig: a genome comparison visualizer. Bioinformatics 27:1009–10.

43. Zimmermann L, Stephens A, Nam SZ, Rau D, Kübler J, Lozajic M, Gabler F, Söding J, Lupas AN, Alva V. 2018. A Completely Reimplemented MPI Bioinformatics Toolkit with a New HHpred Server at its Core. J Mol Biol 430:2237–2243.

44. Larkin MA, Blackshields G, Brown NP, Chenna R, McGettigan PA, McWilliam H, Valentin F, Wallace IM, Wilm A, Lopez R, Thompson JD, Gibson TJ, Higgins DG. 2007. Clustal W and Clustal X version 2.0. Bioinformatics 23:2947–8.

45. Le SQ, Gascuel O. 2008. An improved general amino acid replacement matrix. Mol Biol Evol 25:1307–20.

46. Kumar S, Stecher G, Li M, Knyaz C, Tamura K. 2018. MEGA X: Molecular Evolutionary Genetics Analysis across Computing Platforms. Mol Biol Evol 35:1547–1549.

47. Heidelberg JF, Eisen JA, Nelson WC, Clayton RA, Gwinn ML, Dodson RJ, Haft DH, Hickey EK, Peterson JD, Umayam L, Gill SR, Nelson KE, Read TD, Tettelin H, Richardson D, Ermolaeva MD, Vamathevan J, Bass S, Qin H, Dragoi I, Sellers P, McDonald L, Utterback T, Fleishmann RD, Nierman WC, White O, Salzberg SL, Smith HO, Colwell RR, Mekalanos JJ, Venter JC, Fraser CM. 2000. DNA sequence of both chromosomes of the cholera pathogen Vibrio cholerae. Nature 406:477–83.

48. Chen CY, Wu KM, Chang YC, Chang CH, Tsai HC, Liao TL, Liu YM, Chen HJ, Shen AB, Li JC, Su TL, Shao CP, Lee CT, Hor LI, Tsai SF. 2003. Comparative genome analysis of Vibrio vulnificus, a marine pathogen. Genome Res 13:2577–87.

49. Makino K, Oshima K, Kurokawa K, Yokoyama K, Uda T, Tagomori K, Iijima Y, Najima M, Nakano M, Yamashita A, Kubota Y, Kimura S, Yasunaga T, Honda T, Shinagawa H, Hattori M, Iida T. 2003. Genome sequence of Vibrio parahaemolyticus: a pathogenic mechanism distinct from that of V cholerae. Lancet 361:743–9.

50. Philippe N, Alcaraz JP, Coursange E, Geiselmann J, Schneider D. 2004. Improvement of pCVD442, a suicide plasmid for gene allele exchange in bacteria. Plasmid 51:246–55.

